# The spatial heterogeneity of the gut limits bacteriophage predation leading to the coexistence of antagonist populations of bacteria and their viruses

**DOI:** 10.1101/810705

**Authors:** Marta Lourenço, Lorenzo Chaffringeon, Quentin Lamy-Besnier, Pascal Campagne, Claudia Eberl, Marion Bérard, Bärbel Stecher, Laurent Debarbieux, Luisa De Sordi

## Abstract

Bacteria and their viruses, bacteriophages (phages), are the most abundant components of the mammalian gut microbiota where these two entities coexist over time. The ecological dynamics underlying the coexistence between these two antagonistic populations in the gut are unknown. We challenged a murine synthetic bacterial community with a set of virulent phages, to study the factors allowing phages-bacteria coexistence in the gut. We found that coexistence was neither dependent on an arms race between bacteria and phages, nor on the ability of phages to extend host range. Instead, our data suggest that some phage-inaccessible sites in the mucosa of the ileum serve as a spatial refuge for bacteria, which from there disseminate in the gut lumen. Luminal phages amplify by infecting luminal bacteria maintaining phage throughout the gut. We conclude that the heterogeneous distribution of microbes in the gut contributes to the long-term coexistence of phages with phage-susceptible bacteria. This observation could explain the persistence in the human gut of intestinal phages, such as the crAssphage, as well as the low efficiency of oral phage therapy against enteric pathogens in animal models and clinical trials.

## Introduction

The mammalian gut is a highly complex organ, with anatomical differentiation, lined by a variety of eukaryotic cells types that serve the establishment of a mutualistic relationship between the host and different enteric microbes, including viruses. Bacteriophages (phages) are the most abundant viruses residing in the gut, but their precise role in shaping the microbiome remains unclear (Manrique et al 2017). Changes in the viral and bacterial communities of the gut are increasingly reported to be associated with pathological conditions in humans, including diabetes, inflammatory bowel diseases and colorectal cancer (Hannigan et al 2018b, Manrique et al 2017, Zhao et al 2017). Although fluctuations in the viral communities have been reported in a recent longitudinal study in humans, a large proportion of individual-specific viral contigs remain detectable over time, suggesting that each individual possesses their own viral fingerprint (Manrique et al 2016, Shkoporov et al 2019). However, how phage and bacteria persist together in the gut remain understudied.

Using phage-bacterial model systems, dynamics of the coexistence of predators (or parasites) and preys (or hosts) have been the subject of theoretical and experimental studies, mostly performed *in vitro* and *in silico* (Betts et al 2014, Brockhurst et al 2006, Hannigan et al 2018a, Lenski and Levin 1985, Weitz et al 2013). *In vivo*, the interaction of phages and bacteria in the mammalian gut has been explored in mice and pigs (Galtier et al 2016, Galtier et al 2017, Looft et al 2014, Maura et al 2012a, Maura et al 2012b, Weiss et al 2009). Studies in mice have shown that virulent phages have a limited effect on the targeted bacterial populations within the gut (Galtier et al 2017, Maura et al 2012a, Maura et al 2012b, Weiss et al 2009). Likewise, a large randomized phage therapy trial targeting *Escherichia coli* diarrhea in Bangladeshi children showed no evidence for *in vivo* replication of oral phages in the gut (Sarker et al 2016). Nevertheless, these studies in mice and human showed that virulent phages, despite their limited impact on the host bacterial population, could persist in the mammalian gut for several weeks. A similar situation was recently described in the human gut where the crAssphage coexists with its highly abundant Bacteriodetes bacterial host (Guerin et al 2018, Shkoporov et al 2018, Yutin et al 2018).

Several factors are invoked to understand the coexistence between phages and bacteria, such as arms race dynamics with resistance development to phage infection and viral counter-resistance, the inherent or evolved ability of the phage to infect multiple hosts, and the distribution of these two antagonist populations in structured environments (Brockhurst et al 2006, Doron et al 2018, Galtier et al 2017, Heilmann et al 2012, Hilborn 1975, Labrie et al 2010). Animal models provided data on the segmental distribution of phages and bacteria along the length of the gut, but they lack resolution along the radial dimension (Galtier et al 2017, Maura et al 2012a). In addition, mono-colonized mice lack aspects of competitive and synergistic interspecies interactions of microbes (Weiss et al 2009). As an approach to more realistic conditions, we investigated the contribution of factors involved in the coexistence of phages and bacteria within the gut environment using the synthetic Oligo-Mouse-Microbiota comprising 12 distinct strains (OMM^12^) (Brugiroux et al 2016). This model allows following defined pairs of phages and bacteria in the gut without the need of antibiotics treatment as a confounding factor commonly used to study enteric pathogens *in vivo* (Croswell et al 2009). We established stable colonization of two *E. coli* strains in gnotobiotic OMM^12^ mice and studied the population dynamics of these strains in the presence of virulent phages. We found that phages were less abundant in the mucosal part of ileal sections compared to *E. coli* levels. This observation provides an explanation for the lack for phage-resistant mutant selection and the limited efficacy of virulent phages in reducing intestinal bacterial loads.

## Results

### *E. coli* strains Mt1B1 and 55989 colonize the gut of OMM^12^ mice

Mice harbouring the OMM^12^ consortium (*Acutalibacter muris* KB18, *Akkermansia muciniphila* YL44, *Bacteroides caecimuris* I48, *Bifidobacterium longum subsp. animalis* YL2, *Blautia coccoides* YL58, *Clostridium clostridioforme* YL32, *Clostridium innocuum* I46, *Enterococcus faecalis* KB1, *Flavonifractor plautii* YL31, *Lactobacillus reuteri* I49, *Muribaculum intestinale* YL27 and *Turicimonas muris* YL45) were orally exposed to the murine commensal *E. coli* strain Mt1B1 as a first attempt to test the capacity of an *E. coli* strain to colonize the gut of OMM^12^ mice. Mice became colonized within two to three days, forming a population that remained stable during more than two weeks (Fig. 1A). Mice did not exhibit signs of discomfort or change in feces consistency. Twelve days after inoculation of strain Mt1B1, intestinal sections (ileum, cecum, colon) were examined and the location of strain Mt1B1 was determined by fluorescence *in situ* hybridization (Fig. 1B and Fig. S1). Strain Mt1B1 was found in all sections of the gut, including the ileum, consistent with the location from which it was isolated (mucosa from the ileum of conventional laboratory mice) (Garzetti et al 2018, Lagkouvardos et al 2016). Similar results were obtained with the enteroaggregative *E. coli* strain 55989, a pathobiont, demonstrating the flexibility of the OMM^12^ model to accommodate novel strains (Fig S2). As previously reported in conventional mice, the gut colonization by strain 55989 did likewise not result in any sign of gut disease (Maura et al 2012a).

**Figure 1.**
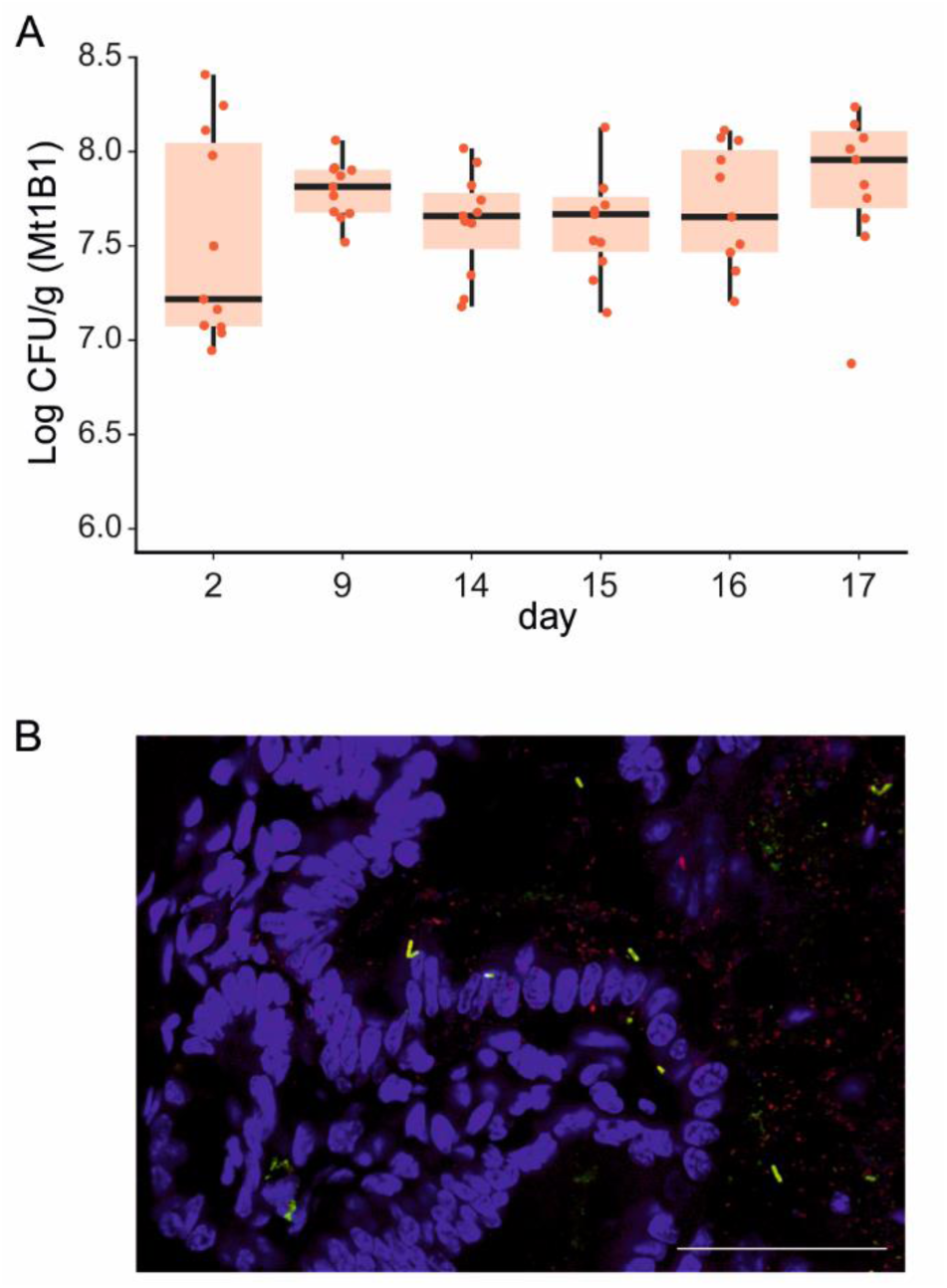
*E. coli* strain Mt1B1 stably colonizes the gut of the OMM^12^ mice. **A.** Fecal levels of *E. coli* strain Mt1B1 at the indicated time points for each OMM^12^ mouse (n=11) receiving a single dose of 10^8^ cfu by oral gavage at day 0. Red dots, individual values; horizontal bar, median; box, 25^th^-75^th^ quantiles, vertical bars, min/max values (within 1.5 x interquartile interval). **B.** Localization by FISH of the strain Mt1B1 in the ileal section of Mt1B1-colonized OMM^12^ mice. Intestinal cells (nuclei) were stained with DAPI (purple), and Mt1B1 (red+green=yellow) and Eubacteria (red) were stained with specific FISH probes. A representative image from a group of four mice is presented (images from colon and cecum are presented in Figure S1). Scale bar, 50µm.

### Isolation and characterization of Mt1B1 phages

We isolated and purified 28 phages infecting strain Mt1B1 from four sources of sewage water. The host-range of these 28 candidates were then characterized, with a panel of 98 different strains of *E. coli*. Three phages (P3, P10, P17) were chosen for this study on the basis of their unique host range (Table S1). All three phages infected strain Mt1B1 in liquid broth. Phages P3 and P10 displayed similar infection patterns, with rapid lysis followed by a moderate bacterial regrowth at 1.5 hours, whereas phage P17 halted the growth of strain Mt1B1 for several hours and slow bacterial regrowth resumed only after more than 10 hours. When used in combination, these three phages caused rapid lysis followed by very slow regrowth (Fig. 2A).

**Figure 2.**
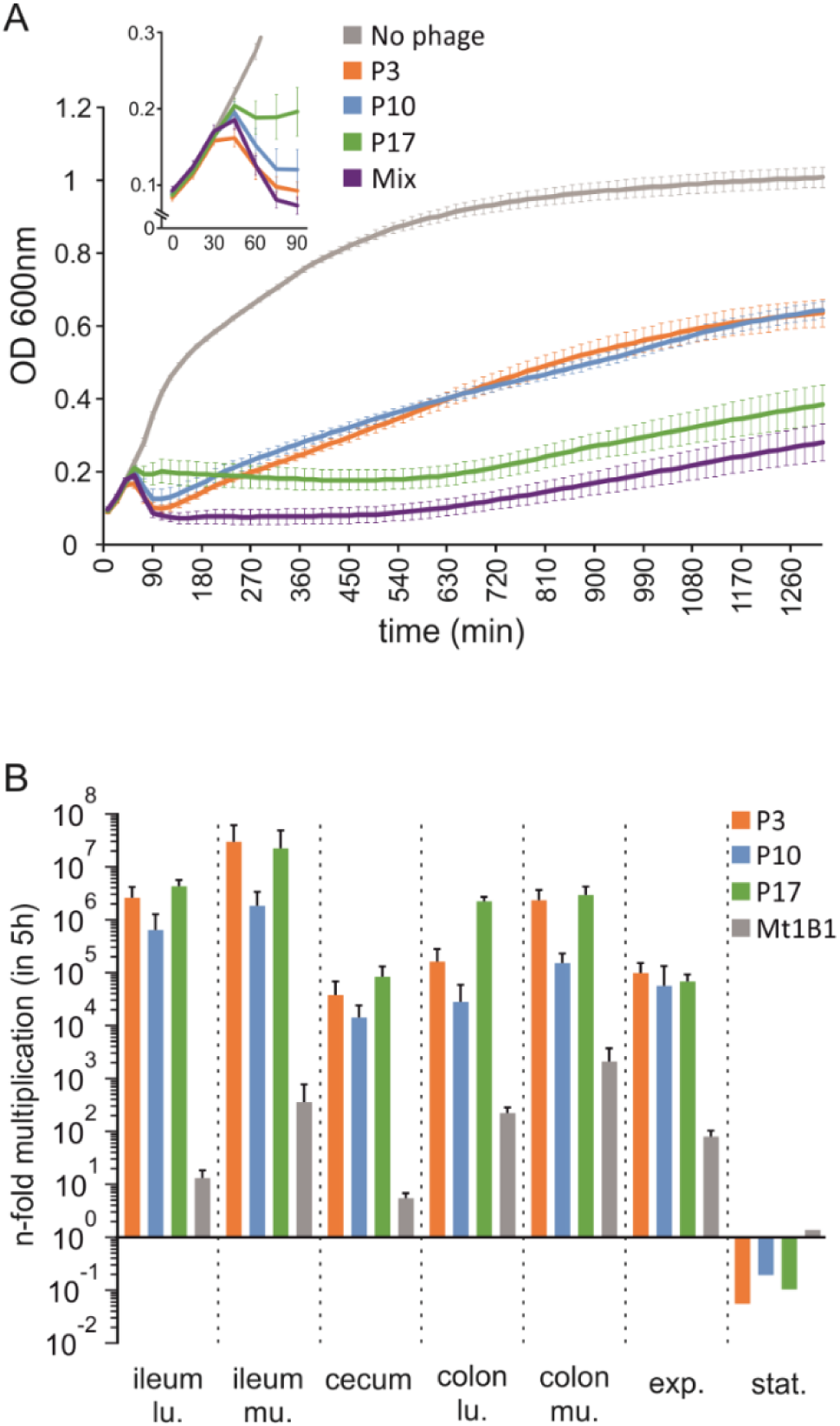
Phages P3, P10 and P17 infect strain Mt1B1 both *in vitro* and *ex-vivo*. **A.** Growth curves for strain Mt1B1 (n=3 for each condition) in liquid broth in the absence (grey) or presence of phage P3 (orange), P10 (blue) or P17 (green), added at t=0, or of a cocktail of these three phages (purple; equal proportions of each) added at t=0. Each phage dose is 100 fold lower than the number of bacteria at t=0. The inset shows an enlargement for early time points. (error bars represent standard error of the mean (SEM) **B.** Amplification over 5h (n=3 biological replicates) of individual phages (P3, orange; P10, blue; P17, green; each at an MOI of 1 x 10^-2^) and Mt1B1 cells (grey) in indicated homogenized gut sections (lu., luminal; mu, mucosal) from Mt1B1-colonized OMM^12^ mice and flasks with Mt1B1 cells in exponential (exp.) (OD_600nm_=0.5) or stationary (stat.; 24h) growth phases. N-fold multiplication relative to the initial number of phages added at t=0, which was approximately 100 fold lower than the amount of Mt1B1 cells, is shown (means with standard deviation).

Phages P3 and P10 had similar affinity constants (5.26×10^-10^±2.8×10^-10^mL.min^-1^ and 4.60×10^-10^±2.15×10^-10^mL.min^-1^, respectively) and adsorption kinetics (90% of phages adsorbed in 2.8 ±0.14 min and 2.03 ±0.19 min, respectively), whereas phage P17 had a stronger affinity for the bacterium (1.83×10^-10^±5.60×10^-11^mL.min^-1^) but slower adsorption kinetics (5.53 ±1.28 min) (Fig. S3). The genomes of these three phages were sequenced and analysed. Phage P3 (40.1 kb; 47 ORFs) belong to Teseptimavirus genus and phage P10 (45.1 kb; 56 ORFs) to Zindervirus genus, both included in the *Autographivirinae* subfamily of *Podoviridae*, whereas phage P17 (150.9 kb; 295 ORFs) is closely related to phage ESCO13 and belongs to an unclassified subfamily of *Myoviridae* (Table S2). The annotation of predicted ORFs revealed an absence of integrases and recombinases homologs, with 57.44%, 75% and 91.53% of ORFs having unknown functions in phages P3, P10 and P17 respectively (Table S3).

We then assessed the capacity of each phage to replicate in gut sections collected from Mt1B1-colonized OMM^12^ mice, in an *ex-vivo* assay previously used to demonstrate the differential activities of phages along the gut (Galtier et al 2017, Maura et al 2012a). We collected samples of the ileum, cecum and colon from the mice. We then separated out the mucosal and luminal parts of the ileum and colonic tissues. We compared the replication of the three phages in these *ex vivo* samples with *in vitro* Mt1B1 planktonic cultures, at exponential and stationary growth phases. Similar patterns were observed for all three phages, with efficient replication in all tested gut sections and in cultures from exponential phase, while no amplification was observed on cells at stationary phase (Fig. 2B). We concluded that Mt1B1 cell growth is necessary for amplification of these three phages.

We then investigated the ability of these three phages to infect *in vivo* the 12 bacteria of the synthetic OMM^12^ community, by administering a single oral dose (6×10^7^ PFU) of a mixture of the three phages (equal amounts of each phage) to OMM^12^ mice not inoculated with strain Mt1B1. Phage levels in feces and in some gut sections (cecum and luminal part of the colon) remained low (about 1×10^4^ pfu/g) 24 and 48 hr after administration. In both the luminal and the mucosal sections of the ileum, and in the mucosal part of the colon, the number of phages was below the threshold of detection (Fig. S4). This finding indicates that none of the Mt1B1 phages multiplied *in vivo* on any of the 12 strains within this time interval. These observations are consistent with findings from the administration of phages to human volunteers or to conventional mice not colonized with the targeted bacteria (Bruttin and Brussow 2005, Maura et al 2012b, Weiss et al 2009).

### A phage cocktail induces a weak but significant reduction in *E. coli* numbers in the gut

We next investigated how Mt1B1-colonized OMM^12^ mice responded to an identical oral three-phage challenge. receiving equal amounts of each of the three phages in a single combined dose of 6×10^7^ PFU would reproduce the long-term coexistence of phages and their target bacterial populations within the gut. OMM^12^ mice were fed strain Mt1B1 once by gavage, and colonization was monitored over nine days. A single dose of the three phages was then administered. Over the following 15 days, the levels of bacteria and phages in the feces were monitored (Fig. S5). These two populations coexisted in similar proportions showing that the OMM^12^ model is suitable to study this coexistence. Since phages replicate *in vivo* but did not exert a strong impact on the fecal levels of strain Mt1B1, we asked whether we could improve this impact by increasing the phage load in repeating the administration of phages during three consecutive days. Such a setting is approaching a phage therapy treatment targeting bacterial pathogens residing in the human gut (Corbellino et al 2019). In two independent experiments, we observed that the group receiving phages displayed on each of the following three days a decrease in fecal levels of strain Mt1B1 compared to the control group of mice receiving no phage (Fig. 3A). These two groups of mice are significantly different when comparing the overall levels of strain Mt1B1 over time (Fig. 3A, p=0.007; TableS4). During the same period, phage levels remained approximately stable (Fig. 3B). In addition, the similar levels of phages and of strain Mt1B1 in fecal samples demonstrate that the coexistence of these antagonistic populations persist despite the repeated application of phages (Fig. 3C).

**Figure 3.**
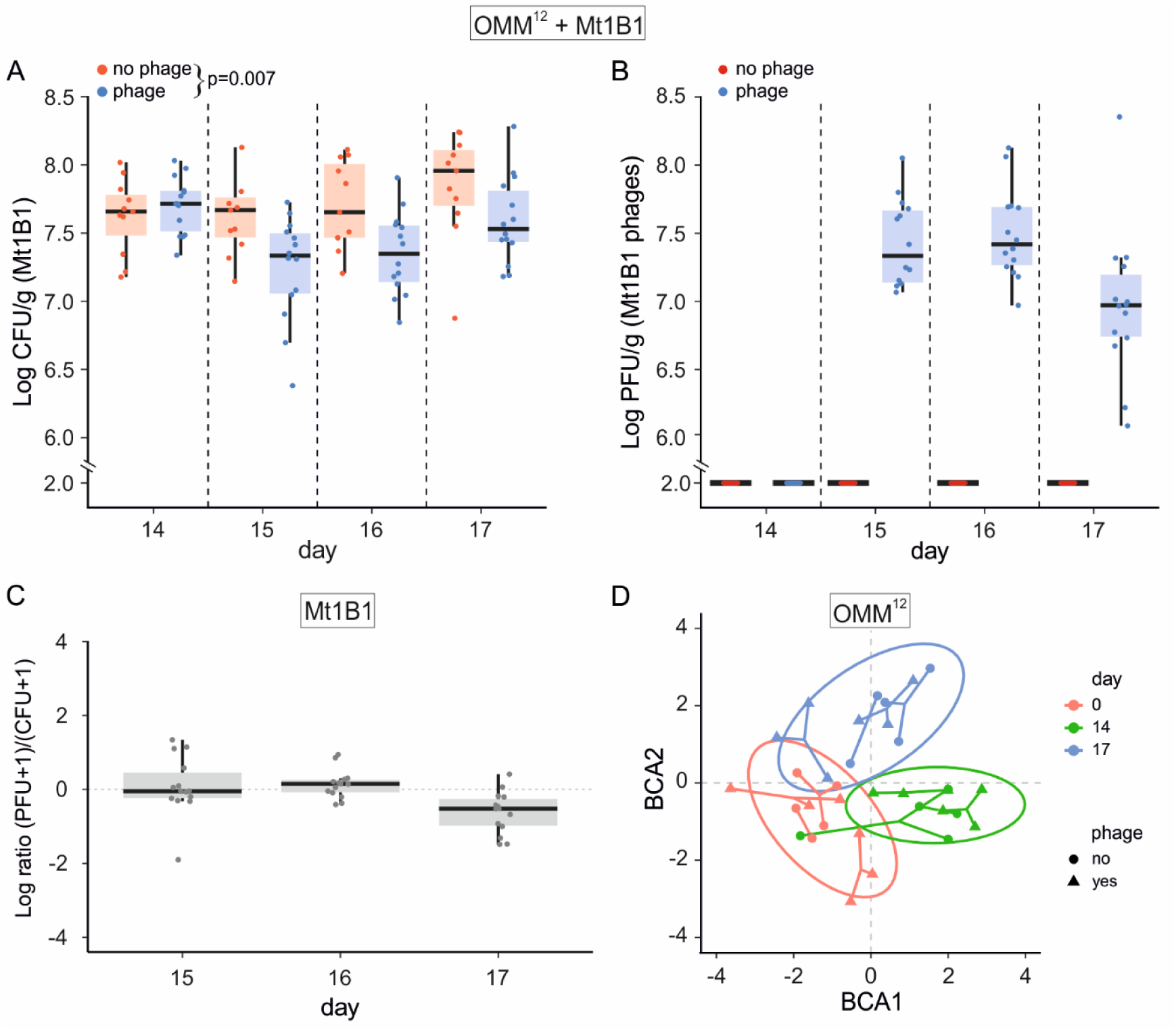
The coexistence of virulent phages with their target strain does not affect the microbiota composition of the OMM^12^ mice. **A.** OMM^12^ mice (n=25) were colonized for 14 days before receiving PBS (red, n=11) or the three phages P3, P10 and P17 together (blue, n=14; 6×10^7^ PFU per dose made of the same amount of each phage) by oral gavage once daily on three consecutive days. Levels of *E. coli* strain Mt1B1 in the feces were recorded over time. **B.** Phage titers from the fecal samples reported in panel A. **C.** Phage:bacteria ratio for fecal samples collected on days 15, 16 and 17 demonstrating the coexistence of the two populations. Dots, individual values; horizontal bar, median; box, 25^th^-75^th^ quantiles, vertical bars, min/max values (within 1.5 x interquartile interval). **D.** Between-group PCA (BCA, axes 1 and 2) for the 16S rRNA qPCR data for mice receiving PBS (n=5, filled circle) or the phage cocktail (n=6, filled triangle) by oral gavage at the indicated time points (see the colors indicated) for 10 bacteria from the microbiota of the OMM^12^ mice (strains YL2 and KB18 were not detected).

### Phage replication does not trigger shifts in OMM^12^ bacterial community

We next assessed the stability of the gut microbiota of both phage treated (n=6) and non-treated (n=5) mice groups by 16S qPCR. This quantification was performed at day0 (before colonization with the Mt1B1 strain), day14 (before the gavage with phage) and day17 (one day after the third phage gavage). Changes in the bacterial community were analysed by performing a between-class principal component analysis (PCA), taking into account days, cages and phage inoculation. Results identified daily fluctuations as the main source of the observed variations regardless of the presence or absence of phages (Fig. 3D, Table S5). This finding is also consistent with the lack of amplification of these phages in OMM^12^ devoid of *E. coli* strain Mt1B1 (Fig. S4A) and suggest that the high fecal phage titer is not the result of the infection of one of the 12 initial strains ruling out a possible off-target amplification. Moreover, results show that the phage replication on one of the bacterial populations present in the gut environment does not significantly affect the stability of the remaining bacterial community, which is consistent with previous reports in the literature (Cieplak et al 2018, Galtier et al 2016). However, a recent report showed that when some phages provokes >2log reduction of the level of their targeted bacterial population, perturbation of the remaining bacterial community is observed in a gnotobiotic mouse model (Hsu et al 2019).

### A unique dose of a single phage is sufficient to initiate its coexistence with its host in the gut

Next, we repeated the experiment in OMM^12^ mice by using *E. coli* strain 55989 and its virulent phage CLB_P2 (Maura et al 2012b). A single dose of phage CLB_P2 (1×10^8^ pfu) did not significantly reduce fecal levels of strain 55989 compared to the control group, while the phage titer remained stable and around 10^7^ pfu/mL within 3 days (Fig S6). Therefore, the coexistence of phages and bacteria in the gut was observed with different pairs of phages and bacteria. Altogether our data suggest that factors limit the impact of phages on *E. coli* strains. One such factors is the selection of phage resistant mutants.

### Maintenance of bacterial populations is not caused by the emergence of phage-resistant clones

To control for the emergence of phage-resistant *E. coli* populations, Mt1B1 clones (n=160, 20 from each of the 8 mice of the short-term experiment; n=120, 24 from each of the 5 mice of the long-term experiment) and 55989 clones (n=75 to 94, from each of the 6 mice) were isolated from fecal samples on the last day of the experiment. All Mt1B1 and 55989 clones were susceptible to each of their respective phages. We also tested for the emergence of phage-resistant clones from several gut sections (luminal and mucosal parts from ileum and colon, together with cecum samples). All of them (n=800; 20 colonies from each organ section of the 8 mice of the short-term experiment with strain Mt1B1; n=75 to 94 colonies from each organ section of the 6 mice with strain 55989) remained susceptible to each of their respective phages. We can therefore conclude that phage-resistant clones did not reach the detection threshold, ruling out a major role of coevolutionary dynamics (arms-race) in the coexistence of virulent phages and bacteria in the gut.

### Spatial heterogeneity of the gut can explain phage-bacteria coexistence

In the absence of evidence for off-target phage amplification or emergence of phage-resistant bacteria, we hypothesized that the spatial heterogeneity of the mammalian gut might create local refuges, where bacteria remained protected against phage predation (Heilmann et al 2012). We tested this hypothesis by examining several gut sections (luminal and mucosal parts from ileum and colon) from mice sacrificed 24 hr after either the last administration of phages in Mt1B1-colonized mice or the third day following a single CLB_P2 administration in 55989-colonized mice. We found that strain Mt1B1 levels from the group of mice receiving phages were significantly lower in all sections, except for the colon luminal sections, compared to the control group (Fig.4A and Table S6). Phage levels were similar in all sections except the mucosal part of the ileum, in which they were strongly reduced (Fig. 4B). A very similar trend was observed from the analysis of gut sections of 55989-colonized mice receiving a single dose of phage CLB_P2 compared to control group. Notably, we found a significant reduction of 55989 levels in the luminal part of the ileum and a lower density of phage CLB_P2 in the mucosal part of the ileum when comparing conditions of presence/absence of CLB_P2 (Fig. 4C,4D).

**Figure 4.**
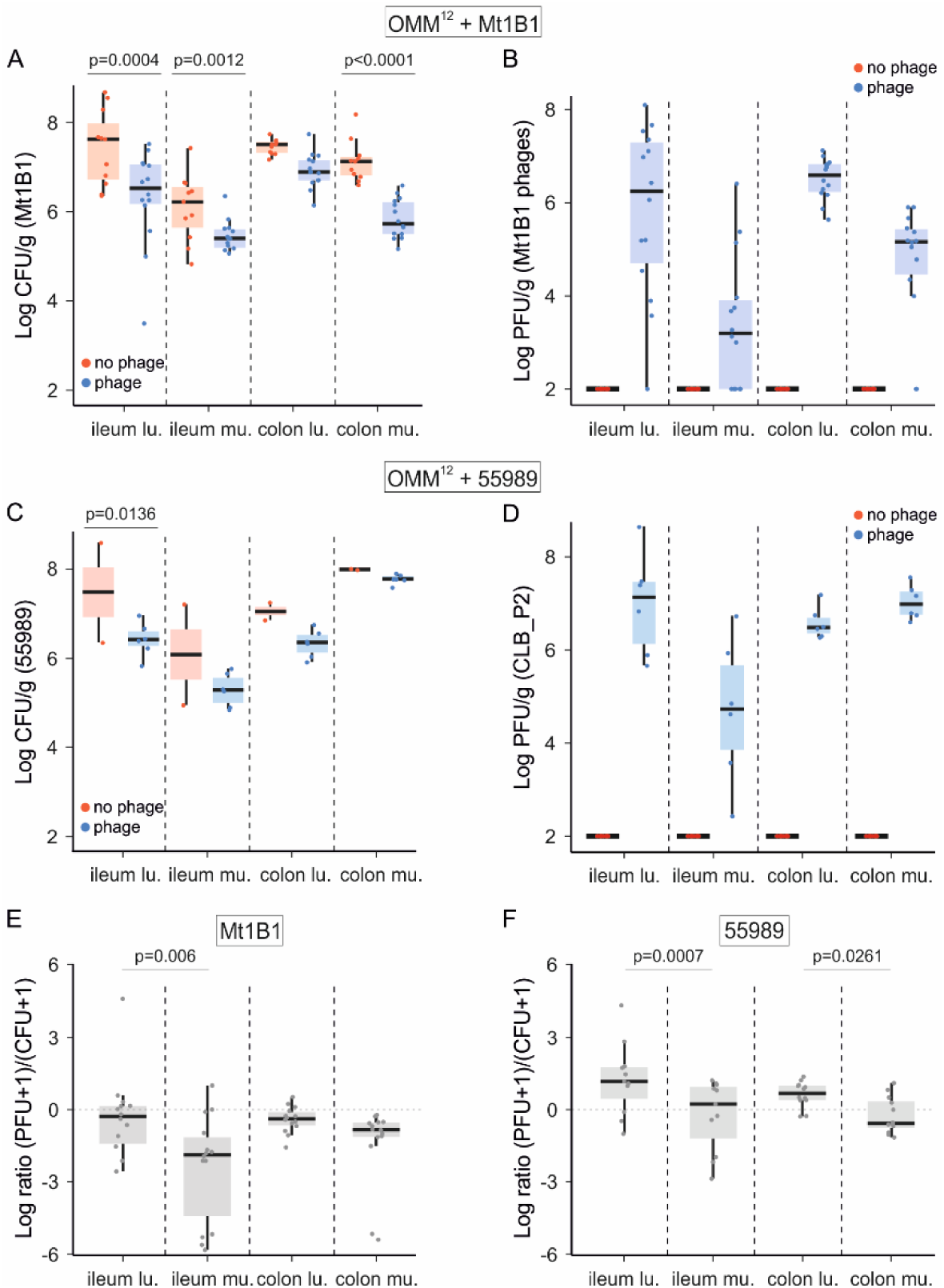
The differential abundance of bacteria and phages in gut sections is consistent with source-sink dynamics. **A.** OMM^12^ mice (n=25) were colonized during 14 days before receiving PBS (red, n=11) or the three phages P3, P10 and P17 together (blue, n=14; 6×10^7^ PFU per dose, composed of equal proportions of each phage) by oral gavage once daily on three consecutive days. Mice were then sacrificed and the abundance of *E. coli* strain Mt1B1 in indicated gut sections was determined. **B.** Phage titers for the samples reported in panel A. **C.** OMM^12^ mice (n=8) were colonized during 7 days before receiving PBS (red, n=2) or a single dose of phage CLB_P2 (blue, n=6; 1×10^8^ PFU) by oral gavage. Mice were sacrificed 3 days later and the abundance of *E. coli* strain 55989 in indicated gut sections was determined. **D.** Phage titers for the samples reported in panel **C**. **E.** Phage:bacteria ratio for the indicated gut sections in panels A and B. **F.** Phage:bacteria ratio for the indicated gut sections in panels **C** and **D.** Dots, individual values; horizontal bar, median; box, 25^th^-75^th^ quantiles, vertical bars, min/max values (within 1.5 x interquartile interval).

The ratio of phages to bacteria in organs from both Mt1B1- and 55989-colonized mice show that the mucosal sections of the ileum exhibit significantly lower phage/bacteria ratios (p=0.006; Table S6 add CLB_P2 data) than the corresponding luminal sections (Fig. 4E, 4F). Thus, the mucosa seems to be less accessible to phages than the lumen and may constitute a refuge in which a reservoir of the *E. coli* population is protected against the phage predation. Phage diffusion in the gut may be limited by mucins and other glycoproteins, lipids and DNA molecules (Johansson et al 2011, Qi et al 2017). A specific *in silico* search for immunoglobulin motifs, which have been shown to favour phage binding to mucus, revealed that only ORF 118 of phage P17 possess such a motif (homologous to the bacterial Ig-like domain (Big2)) (Barr et al 2013, Fraser et al 2006). The absence of these motifs in three out of the four phages studied here is consistent with our *in vivo* observation that these phages are less abundant in the mucosal sections.

## Discussion

The intestinal microbiota of mammals is relatively stable over time in healthy subjects while it contains large antagonistic populations of bacteria and phages. The mechanisms that underlie this apparently peaceful coexistence are unknown. Several hypotheses can be drawn from years of *in vitro* studies of isolated systems (one phage/one bacterium) or more recently from environmental studies (Buckling and Rainey 2002, Fortuna et al 2019, Horne 1970, Laanto et al 2017, Meyer et al 2016). However, none of them has yet been tested in relevant animal models. Using a gnotobiotic mouse model, two *E. coli* strains and four different virulent phages, we show that the emergence of phage-resistant bacteria and the possibility to infect off-target bacteria can be excluded as major factors supporting coexistence. Instead, we found that the spatial distribution of phages and bacteria along the radial axis of the gut fits the classical ecological theory of source-sink dynamics (Holt 1985). Our data suggest that bacterial refuges in the mucosal layer serve as a source, with the lumen acting as a sink in which the phages can infect their target. As phages do not exert a major direct selective pressure on the source, the bacteria reaching the gut lumen remain susceptible enabling the phages to maintain their density in the lumen over time (Fig 5).

**Figure 5.**
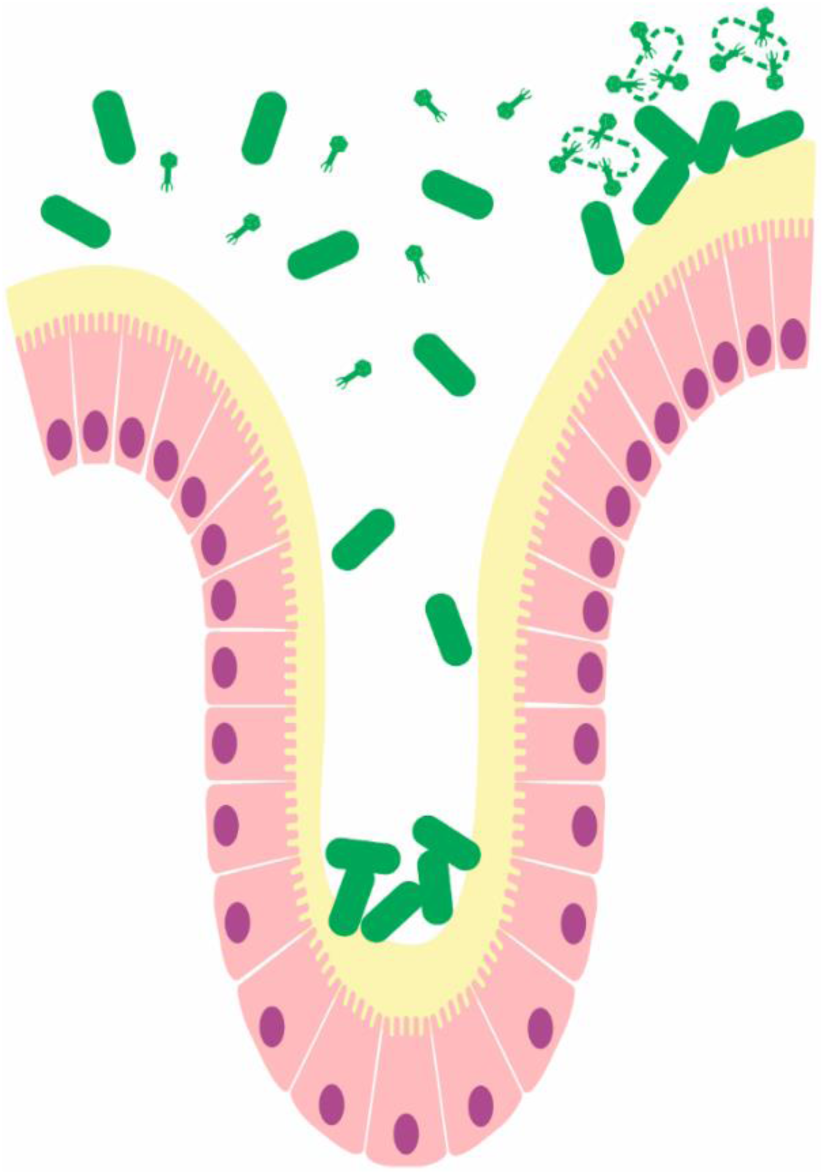
Schematic representation of the source-sink dynamics between bacteria and virulent phage populations in the gut. Bacteria located close to the mucosal layer (in yellow) of epithelial cell form refuges that phages cannot reached. In the intestinal lumen phages encounter enough bacteria to persist by killing bacteria (dotted lines).

In phage/bacterium interaction studies, the prey-predator dynamics have been classically studied through the prism of phage-resistant bacteria and counter resistant phages including phage and bacteria densities. In our experimental conditions, we found that our data fit with a source-sink model. Recently, an *in vitro* study has proposed the model named “leaky resistance” to explain the maintenance of virulent phages by the high rate of transitions between phage-resistant and phage-susceptible bacteria within the populations (Chaudhry et al 2018, Silveira and Rohwer 2016). However, in our experimental data collected from two couples of phages and bacteria we failed to isolate phage-resistant bacteria within the entire gut. This suggests that in our conditions, including a single or repeated phage input, these phage-resistant bacteria may either carry a high cost or have a low selective value (Gomez and Buckling 2011, Koskella et al 2012). Coupled with the concept of costly phage resistance mutations, our results are in agreement with observations from *in silico* individual based stochastic spatial models, which showed that structured environments can create spatial refuges that lead to coexistence between bacteria and phages without the emergence of resistant clones (Heilmann et al 2012, Sousa and Rocha 2019). In addition, the lower abundance of phages in the ileum mucosa could be caused by the reduced bacterial density at this site, compared to the other gut sections, according to a density-dependent phenomenon that was previously observed in several *in vitro* structured environments (Eriksen et al 2018). This density-dependent phenomenon has been linked to the “proliferation threshold” that proposes that a minimum number of bacteria is needed for the phage to initiate infection and amplify (Payne et al 2000, Wiggins and Alexander 1985). Note that the source-sink and the “proliferation threshold” hypotheses are not mutually exclusive to support the presence of bacterial refuges driving the long-term coexistence of phages and susceptible bacteria.

The presence of several IgG-like motives in capsid proteins of phages mediates their interaction with mucus layers (Barr et al 2013). The analysis of the four phage genomes to detect IgG-like motives revealed that only one contains a predicted protein with a single motif with a weak score. Therefore, the lack of such motives is in agreement with our data showing that the density of phages in the mucosal sections of both ileum and colon is significantly lower than in the corresponding luminal sections. We hypothesize that the gut displays a mucosal-luminal gradient of phage density. Consequently, bacterial populations are exposed to different phage concentrations.

Several other processes, not directly tackled in this study like the influence of abiotic factors, can contribute to the coexistence of phage and bacteria populations in the mammalian gut (Lourenco et al 2018, Scanlan et al 2019). In particular, phenotypic resistance, described as the ability of bacteria to oscillate between phage-susceptible and phage-resistant cells by modulating the expression of genes that encode for phage receptors, led by environmental or cellular stochasticity (Bull et al 2014, Chapman-McQuiston and Wu 2008).

Our data show that the gnotobiotic OMM^12^ model is particularly well suited to study the molecular mechanisms involved in phage bacteria dynamics in the gut. OMM^12^ community establishes a long-term stable composition over several mouse generations and can be used as platform to flexibly incorporate additional bacterial strains to the community (Brugiroux et al 2016, Herp et al 2019, Studer et al 2016). Interestingly, Hsu and colleagues recently reported the use of similar synthetic microbiota models to study phage bacteria interactions with two major differences. First, axenic mice were inoculated with bacterial strains during only two weeks before the inoculation of phages, compared to the permanently colonized OMM^12^ mice. Therefore, the immune system is likely stimulated by microbial components for the first time along the course of their experiments. Second, the 10 bacterial strains chosen are from human origin, instead of mouse origin for OMM^12^, and may rapidly undergo genetic adaptation to the mouse environment (Barroso-Batista et al 2014, Lourenco et al 2016). Despite these differences, both mouse models confirm that the inoculation of virulent phages in the gut generally leads to the coexistence of phages and bacteria. Hsu et al. found that *Enterococcus faecalis* phage resistant mutants increased over time, while we could not detect such events. In addition, they found that some phages have strong impact on abundance of cognate host bacteria, while such impact was not previously observed in several mouse models (Maura et al 2012a, Weiss et al 2009). This suggests that different phage-bacteria couples may exhibit distinct eco-evolutionary dynamics which points at even higher complexity of studying these interactions in natural environments (Shkoporov and Hill 2019).

Additional studies on phage bacteria interactions in the gut to improve our understanding of the infection dynamics are required to develop efficient phage-guided strategies and ultimately obtain firm clinical evidences that are still lacking (Brussow 2017, Sarker et al 2016). A recent case report showed that the *in vitro* isolation of a single virulent phage, used to target one multidrug resistant strain of *Klebsiella pneumoniae*, led to the eradication of this pathogen from the patient’s gut (Corbellino et al 2019). While lacking mechanistic insights it confirms the medical potential of phages to selectively eliminate bacteria residing in the gut.

## Material and Methods

### Ethics statement

All animal experiments were approved by the committee on animal experimentation of the Institut Pasteur and by the French Ministry of Research. C57Bl/6J mice (seven to nine-week-old) OMM^12^ were bred at Institut Pasteur (Paris, France). A total of 50 OMM^12^ mice were used.

### Phage isolation

First, sewage water from four locations was filtered at 0.45μm and mixed with an equal volume of 2X Luria-Bertani (LB) medium. Second, these four mixtures were inoculated with a fresh growing culture of Mt1B1 (OD of 0.4 at 600nm, final dilution 1/200) and incubated on a shaker at 37°C overnight. The next day chloroform (1/10 volume) was added to the flasks and incubated at room temperature for one hour before to be centrifuged at 8000g for 10 min. One mL of the supernatant was mixed with 1/10 vol. of chloroform and centrifuged at 8000g for 5 min. A 100-fold dilution in TN buffer (10mM Tris HCl pH7.5, NaCl 100mM) of the aqueous phase was prepared. 10µL of the undiluted and diluted solutions were spread with an inoculation loop on the top of two separate LB agar plates and allowed to dry for 30 min under a safety cabinet. Subsequently, 1 mL of an exponentially growing culture of Mt1B1 was applied to cover entire each plate; the excess liquid culture was removed and the plates were incubated at 37°C overnight. The next day, individual plaques were picked and resuspended in tubes containing 200μl TN buffer. 1/10 vol. chloroform was mixed and tubes were centrifuged at 8000g for 5 min. These steps of plaque purification were performed three times. Finally, 10µL of the last resuspended plaque were added to 1mL of a liquid culture of Mt1B1 (OD of 0.4 at 600nm) and incubated at 37°C in a shaker for 5 hours. 1/10 vol. of chloroform was mixed and after centrifugation at 8000g for 5 min this stock was stored at 4°C and served as starting solution for large scale lysates.

### Bacterial strains, and host range tests

Bacterial strains including Mt1B1 (DSM-28618) and 55989 are listed in Table S1. Strain Mt1B1-DsRed, was obtained by targeted recombineering using strain LF82-DsRed as template (De Sordi et al 2017). Host range tests were performed as follows: 3µL of PBS-diluted phage solutions (0,2µm filtered sterilized crude lysates adjusted to 10^7^ pfu/mL) were deposited side by side on the lawn of each tested bacterium on agar LB plates. Plates were incubated at 37°C overnight. Phages were grouped according to their host range and three representative phages of the main groups were chosen (Table S1).

### Adsorption assays and phage growth

Three independent adsorption assays were performed for each phage according to the protocol previously described (Chevallereau et al 2016). Data could be approximated using an exponential function and adsorption times were defined as the time required to reach a threshold of 10% of non-adsorbed phage particles. To record phage growth and bacteria lysis, an overnight culture of strain Mt1B1 was diluted in LB broth and grown to an OD_600nm_ of 0.2 from which 150 µL were distributed into each of the wells of a 96-well plate (Microtest 96 plates, Falcon). 10 µL of sterile phage lysates diluted in PBS to obtain a multiplicity of infection (MOI) of 1 x 10^-2^ in each well. Plates were incubated in a microplate reader at 37°C, with a shaking step of 30 sec before the automatic recording of OD_600nm_ every 15 min over 20 hr (Glomax MultiDetection System, Promega, USA).

### Phage genomes sequencing and analysis

Sterile phage lysates were treated by DNase (120 U) and RNase (240 µg/mL) and incubated for 30 minutes at 37°C. RNase/DNase reaction was stopped by adding EDTA (20 mM). Lysates were treated with proteinase K (100 µg/mL) and SDS (0.5%) and incubated at 55°C for 30 minutes. DNA was extracted by a phenol-chloroform protocol modified from Pickard (Pickard 2009). Sequencing was performed using Illumina technology (Illumina Inc., San Diego, CA) MiSeq Nano with paired-end reads of 250bp. Quality of reads was visualised by FastQC v0.10.1 Brabraham Bioinformatics (http://www.bioinformatics.babraham.ac.uk/projects/fastqc/). Assembly was performed using a workflow implemented in Galaxy-Institut Pasteur using clc_assembler v4.4.2 and clc_mapper v4.4.2 (CLC Bio, Qiagen). Phage termini were determined by PhageTerm (Garneau et al 2017) and annotations were performed by the RAST v2.0 server (Aziz et al 2008).

### Search for Ig-like domains on phage genomes

Protein sequences of the four phages were scanned for homologs on Pfam database using the HMMER website (Potter et al 2018). The results were compared to a comprehensive HMM (Hidden Markov Models) database of Ig-like domains found on Pfam that was kindly provided by Dr. Sean Benler.

### *Ex vivo* phage replication assay

Oligo-MM12 mice received 200µL of strain Mt1B1 (10^7^ CFU prepared from an overnight culture in LB at 37°C) in sterile sucrose sodium bicarbonate solution (20% sucrose and 2.6% sodium bicarbonate, pH 8) by oral gavage and three days after were sacrificed to collect and weight intestinal sections (ileum, cecum and colon). PBS was added to each sample (1.75mL for ileum and colon and 5mL for the cecum) before homogenization (Oligo-Macs, Milteny Biotech). A volume of 150µL of each homogenized sample was distributed in the wells of a 96-well plate and 10µL of each individual phage were added to reach an MOI of 1 x 10^-2^, and the plate was incubated at 37°C. A fraction of the homogenized samples was also serially diluted in PBS and plated on Drigalski medium to count Mt1B1 colonies at t=0. Following five hours of incubation, samples were serially diluted in PBS and plated on Drigalski medium as well as on LB agar plates overlayed with strain Mt1B1. Both set of plates were incubated at 37°C overnight. The same procedure was followed for *in vitro* growth assays with bacteria taken during exponential (OD 0.5) or stationary (24 hr) growth phase at 37°C with shaking.

### Murine model of *E. coli* colonization and quantification of phages and bacteria

The long-term coexistence experiment with strain Mt1B1 included 7 mice (5 that received phages and 2 that did not) and lasted 23 days. At day 0 mice feces were collected prior to Mt1B1 administration by oral gavage (200 µL of bacteria resuspended in sodium bicarbonate buffer from an overnight liquid culture). Fecal pellets were transferred in pre-weighted, sterile, 2 ml tubes, weighted and resuspended in 1 ml of PBS. Serial dilutions in PBS were performed and plated onto Drigalsky plates. The three phages (2×10^7^ PFU of each phage in 200 µL of PBS) were administered altogether once by oral gavage at day 9. The level of phages was assessed from serial dilutions in PBS spotted on LB plates overlayed with strain Mt1B1.

Two shorter independent experiments with stronger phage selective pressure were performed with 11 (6 with phages and 5 without) and 14 (8 with phages and 6 without) Mt1B1-colonized mice respectively. At day 0 mice feces were collected prior to Mt1B1 administration (as described above) by oral gavage. Fecal pellets were transferred in pre-weighted, sterile, 2 ml tubes, weighted and resuspended in 1 ml of PBS. Serial dilutions in PBS were performed and plated onto Drigalsky plates. The three phages (2×10^7^ PFU of each phage in 200 µL of PBS) were administered altogether once by oral gavage at day 14, 15 and 16. The level of phages was assessed from serial dilutions in PBS spotted on LB plates overlayed with strain Mt1B1. Each mouse was sacrificed at day 17 to collect feces and intestinal sections all homogenized in PBS using gentleMACS^TM^ OCtoDissociator (Miltenyi Biotec) and plated on both Drigalski plates and LB plates overlayed with strain Mt1B1. The luminal sections correspond to the gut contents that were recovered by pressing the surface of the tissues with the back of a scalpel. The remaining tissues correspond to the mucosal sections that were homogenized together with the organ epithelium.

For experiments with *E. coli* strain 55989 fecal samples were collected on day 1, 3, 7, 8, 9 and 10. On the day 0, after collection of the fecal samples, we proceeded to the administration by oral gavage of *E.coli* strain 55989 the same way than with strain Mt1B1. Approximately 2×10^9^ CFUs/g per mouse, resuspended in a sucrose bicarbonate solution (200g.L^-1^ sucrose, 26g.L^-1^ sodium bicarbonate, pH=8), were administered. At day 7, after collection of the fecal sample, 200µL of either a phage suspension containing approximately 2×10^8^ pfus/mL of phage CLB_P2 (n=12) or PBS (n=6) were administered to the mice by oral gavage. At days 8 and 10, feces were collected before sacrificing and dissecting the mice in order to collect organs. Caecum, ileum and colon were collected, and for the latter two the adherent and luminal part were separated and manipulated as reported above. The levels of bacteria and phages were assessed as reported for the strain Mt1B1.

### Identification of resistant clones

For the experiments using Mt1B1 and the cocktail, 20 clones from each gut section and fecal samples (per mice) were streaked vertically in LB agar plates and subsequently each of the three phages was horizontally streaked across. Plates were incubated at 37°C and phenotype was checked after 5h and overnight.

For the experiments using 55989 and phage CLB_P2, the 75 to 94 isolated colonies per sample were randomly chosen by a robot (Qpix 420; Molecular Devices, Sunnyvale, USA) and cultured overnight in 150µL of LB Lennox in 96 well microplates. The next day, 10µL were added to a new plate filled with 140µL of LB and incubated at 37°C for 2h. Then, 8µL of each clone were spotted on two separated agar plates (LB Lennox) and let dry for 20 minutes. Then, a spot of 4µL of phage CLB_P2 (1×10^5^ PFU/mL for one plate and 1×10^7^ PFU/mL for the second plate) were spotted on top of each bacterial spot. Susceptible clones are defined by the full or partial clearance of the bacterial spot at one or both phage concentrations, while clones not affected by the phage at any concentration are defined as resistant.

### Fluorescence in situ hybridization

Fluorescence in situ hybridization was performed as previously described on intestinal samples from *E. coli* Mt1B1 colonized OMM^12^ mice bred at the LMU Munich where the fecal level of strain Mt1B1 reached 10^9^ CFU/g (Brugiroux et al 2016). Ileal, cecal and colonic tissue was fixed in 4% paraformaldehyde (4°C overnight), washed in 20% sucrose (4°C overnight), embedded in O.C.T (Sakura), and flash frozen in liquid nitrogen. FISH was performed on 7µm sections, using double 3′ and 5′-labelled 16S rRNA targeted probes specific for Enterobacteriaceae (Ent186-2xCy3 (CCC CCW CTT TGG TCT TGC)) and Eubacteria (1:1 mix of Eub338I-2xCy5 (GCT GCC TCC CGT AGG AGT) and Eub338III-2xCy5 (GCT GCC ACC CGT AGG TGT)). 1 μg/mL–1 4′,6-diamidino-2-phenylindole (Roth) was used for DNA staining. Images were recorded with a Leica TCS SP5 confocal microscope (Leica, Wetzlar).

### qPCR for 16S quantification

From homogenized fecal samples, 500 µL were centrifuged at 8.000g for 10 min and the supernatant removed. Pellets were diluted in 500 µL of lysis buffer (500 mM NaCl, 50 mM Tris-HCl, pH 8.0, 50 mM EDTA, 4% sodium dodecyl sulfate (SDS)) and incubated for 15 min at 50°C (Yu and Morrison 2004). Then, 100 µL of lysozyme (25mg/ml) was added and samples were incubated at 37°C for 2 hr. DNA extraction was performed using the Maxwell Cell tissue kit (Promega). The primers, probes and qPCR protocol were used in conformity with previously described methods (Brugiroux et al 2016) with the exception of the SsoAdvancedTM Universal Probes Supermix (BioRad). The qPCR reactions were performed in duplicate and in two independent runs using MasterCycler realplex4 from Eppendorf. Statistical analysis is described in the section below.

### Statistical analysis

Statistical analysis on the number of bacteria and phages were carried out using the lme4 and lmerTest packages of R (Bates et al 2015, Kuznetsova et al 2017). Both CFU and PFU were log10- transformed prior to analysis. In each experiment, two groups of mice were considered, a group exposed to phages and a control (unexposed) group. Beyond these groups, the effects of phages could be assessed based on the abundance of phages (log-PFU). Given the non-linearity of responses, the day at which a measure was performed was considered as a categorical variable. Linear mixed-models were used to account for random experimental effects (i.e., both individual and cage effects). Overall effects were assessed with Analysis of Variance (ANOVA) and post-hoc Tukey’s comparisons and were performed using the *lsmeans* R package (Lenth 2016). 16S-quantification data were analysed using multivariate analysis after standard normalization. A principal component analysis (PCA) was performed with the R package ade4 on the matrix of ΔCt values of 10 bacterial strains (strains YL2 and KB18 were not detected). In addition, a between-group PCA was done in order to assess experimental effects, based on 12 groups of observations: 3 days (0, 14, and 17) and 4 cages (2 exposed, 2 unexposed).

## Author contribution

LD, LDS and ML conceived the study. CE, LC, LDS, ML and QLB performed experiments. LC, ML and PC performed statistical analysis. BS, LD and MB secured funding. LD, LDS and ML drafted the paper and all the authors contributed and approved the submitted version.

## Acknowledgements

We thank Jorge Moura de Sousa and Harald Brüssow for critically reading the manuscript, Dwayne Roach and Anne Chevallereau for valuable discussion. We thank Sean Benler for kindly sharing the comprehensive HMM database of Ig-like domains identified on Pfam database. We thank the members of the Centre for Gnotobiology Platform of the Institut Pasteur (Thierry Angélique, Eddie Maranghi, Martine Jacob and Marisa Gabriela Lopez Dieguez) for their help with the animal work. ML is part of the Pasteur - Paris University (PPU) International PhD Program. ML is funded by Institut Carnot Pasteur Maladie Infectieuse (ANR 11-CARN 017-01). LDS is founded by a Roux-Cantarini fellowship from the Institut Pasteur (Paris, France). QLB is funded by Ecole Normale Supérieure. BS is supported by the German Center of Infection Research (DZIF), the Center for Gastrointestinal Microbiome Research (CEGIMIR), the DFG Priority Programme SPP1656 (STE 1971/4-2 and STE 1971/6-1) and the Collaborative Research Center CRC 1371.

