## Supplemental figures and tables for "The spatial heterogeneity of the gut limits bacteriophage predation leading to the coexistence of antagonist populations of bacteria and their viruses"

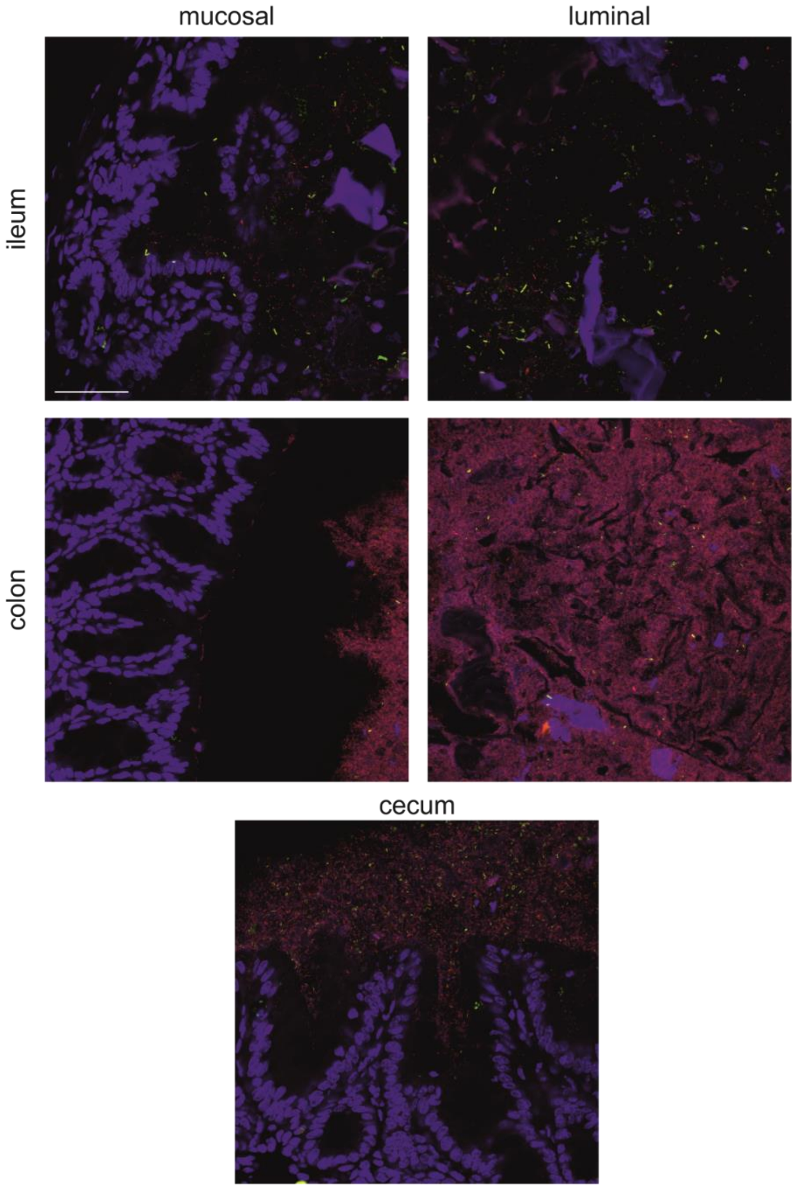


**Figure S1. Localization by FISH of strain Mt1B1 in gut sections from OMM$^{12}$ mice.**

Intestinal cells (nuclei) were stained with DAPI, and Mt1B1 (red+green=yellow) and Eubacteria (red) were stained with specific FISH probes. Representative images from a group of five mice are presented. Scale bar, 50µm.

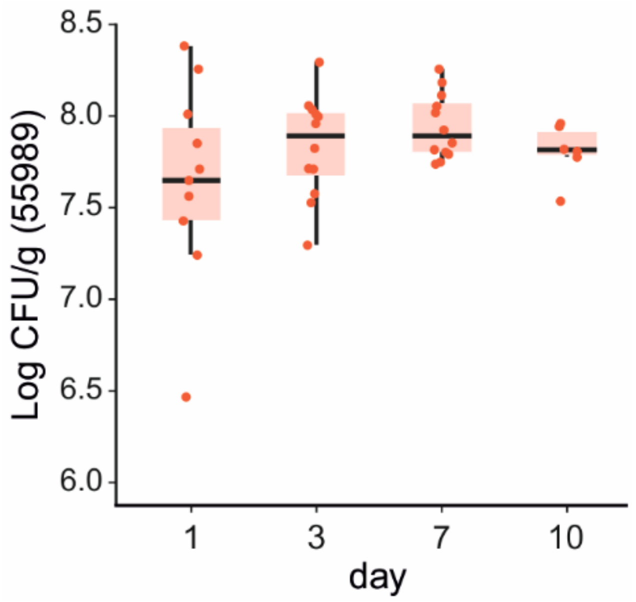


**Figure S2.** ***E. coli*** **strain 55989 can stably colonize the gut of OMM$^{12}$ mice.**

Fecal levels of *E. coli* strain 55989 at the indicated time points for each OMM$^{12}$ mouse (n=10) receiving a single dose of $10^8$ cfu by oral gavage at day 0. Red dots, individual values; horizontal bar, median; box, 25$^{th}$-75$^{th}$ quantiles, vertical bars, min/max values (within 1.5 x interquartile interval)

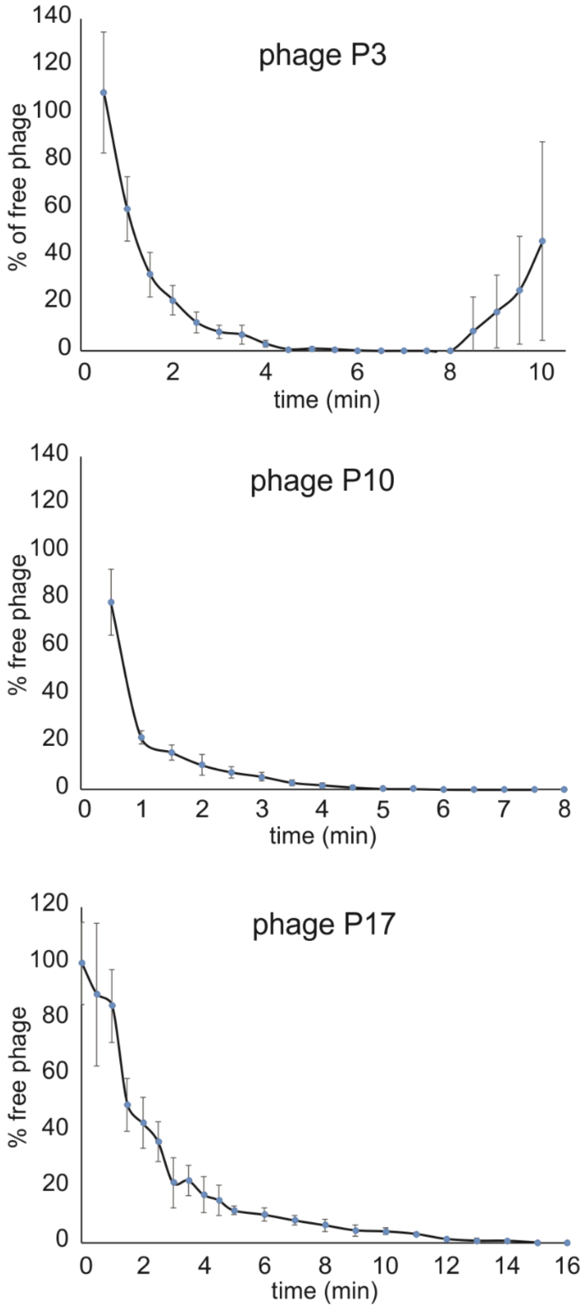


**Figure S3. Mt1B1 phages P3, P10 and P17 have different adsorption rates.**

An exponentially growing culture of strain Mt1B1 in LB medium was infected independently by phages P3, P10 or P17 (MOI= 1 x $10^{-2}$) and samples were collected over time for counting non-adsorbed phages. For phages P3 and P10, samples were taken every 30 s over a period of 10 minutes. For phage P17, sampling was performed every 30 s until 5 minutes, and then every minute until 15 minutes. The percentage of non-adsorbed phages is plotted against time (mean with standard deviations; n=3).

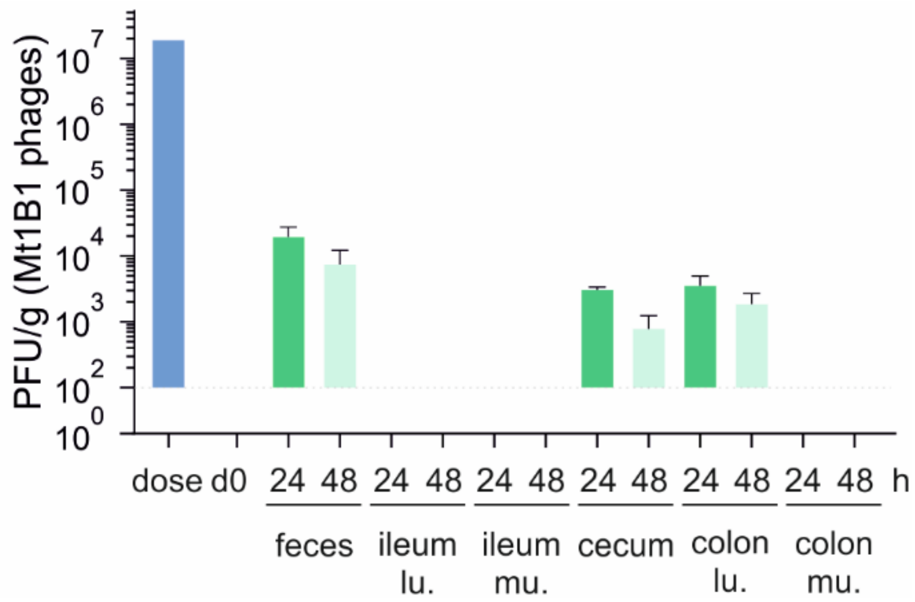


**Figure S4. Phages do not replicate in the gut in the absence of strain Mt1B1**.

Phage titers were assessed from fecal samples and samples from the different sections of the gut collected at 24H (dark green) and 48H (light green) from OMM[12] mice that had received a single dose of the three phages ($3x10^7$ pfu (blue bar); phages mixed in equal proportions). The mean with standard deviations of two independent experiments each with n=2 mice is represented).

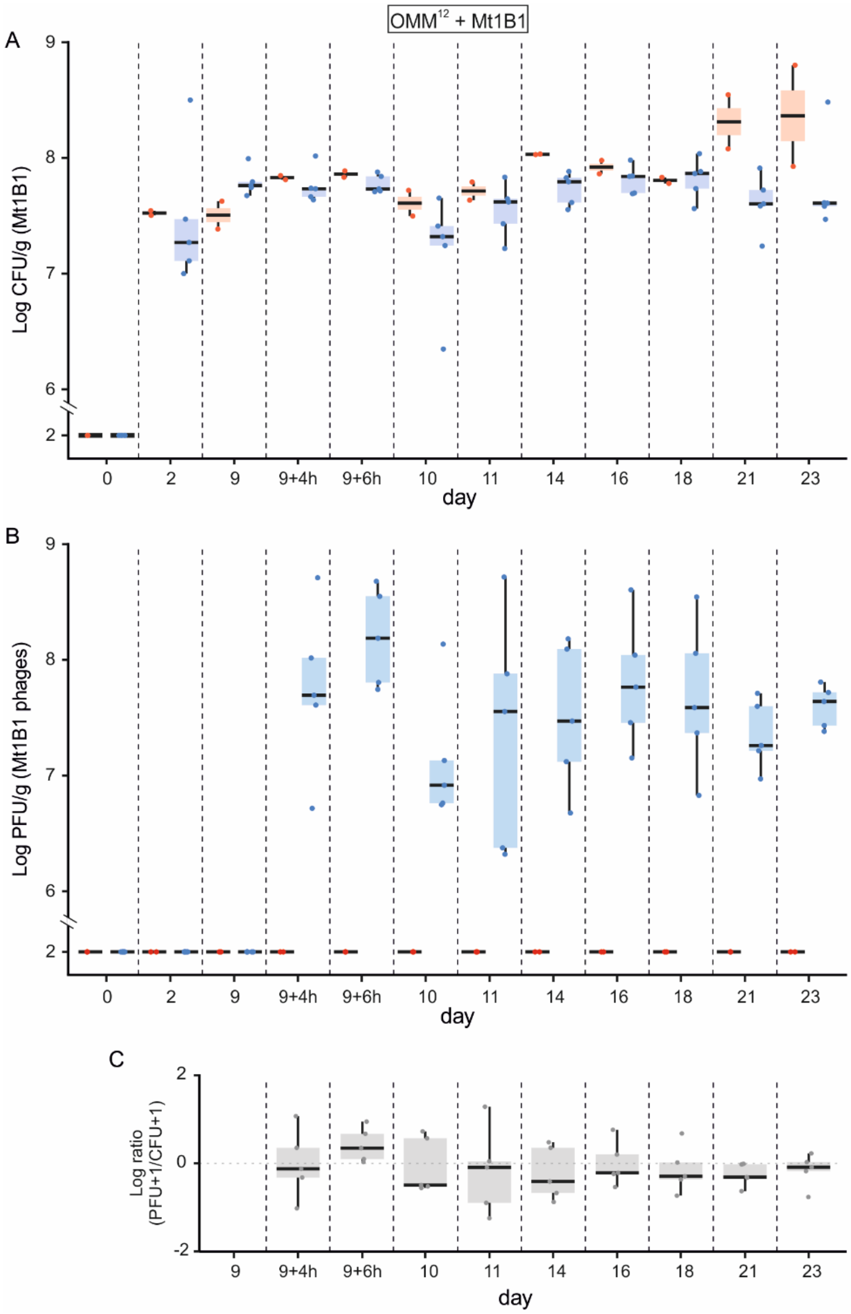


**Figure S5. Mt1B1-colonized OMM$^{12}$ mice are suitable for use in studies of the long-term coexistence of phage and bacteria in the mammalian gut.**

OMM$^{12}$ mice (n=7) were colonized during 14 days before a single administration of PBS (red, n=2) or the three phages P3, P10 and P17 together (blue, n=5; 6x10$^{7}$ PFU per dose, consisting of equal proportions of each phage) by oral gavage on day 9. A. The levels of *E. coli* strain Mt1B1and B. the levels of phages were recorded at the indicated time points. C. Representation of the ratios phage:bacteria over time showing the stability of the coexistence. Dots, individual values; horizontal bar, median; box, 25$^{th}$-75$^{th}$ quantiles, vertical bars, min/max values (within 1.5 x interquartile interval).

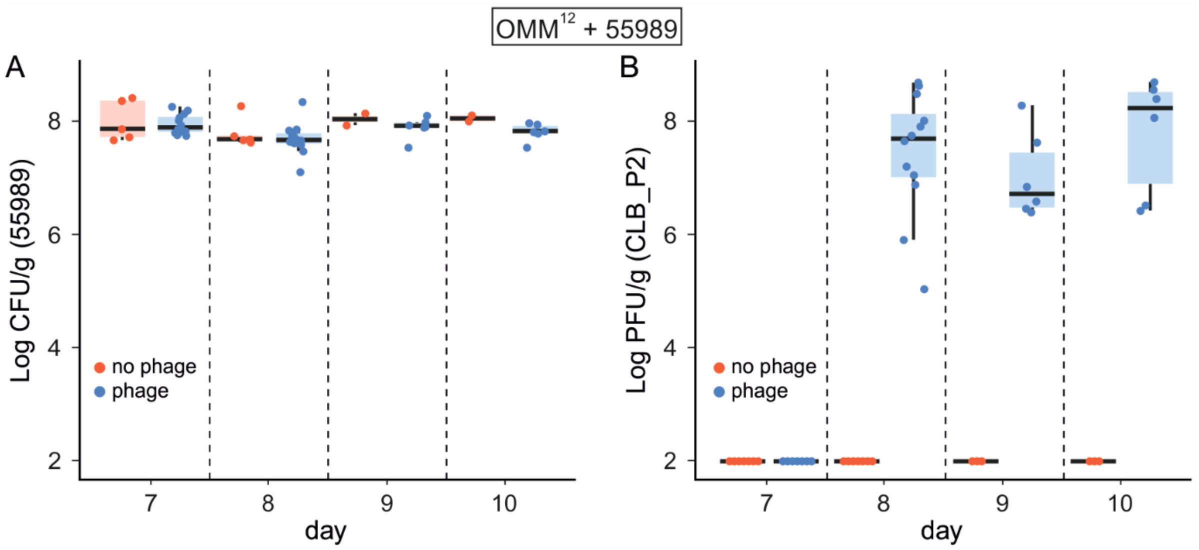


**Figure S6. A single dose of the phage CLB_P2 is sufficient to establish the coexistence of phages and bacteria in the mammalian gut.**

OMM$^{12}$ mice (n=17) were colonized during 7 days before a single administration of PBS (red, n=5) or the phage CLB_P2 (blue, n=12; 1x$10^{8}$ PFU) by oral gavage on day 9. The levels of *E. coli* strain 55989 were recorded at the indicated time points. Dots, individual values; horizontal bar, median; box, 25$^{th}$-75$^{th}$ quantiles, vertical bars, min/max values (within 1.5 x interquartile interval).

**Supplemental Tables**

**Table S1. Host range of the 28 candidate phages isolated for strain Mt1B1.**

(see separated Excel file)

**Table S2. List of the closest phage homologs to phages P3, P10 and P17.**

(see below)

**Table S3. Table of genome annotations for phages P3, P10 and P17.**

(see separated Excel file)

**Table S4. Statistical analysis, with a mixed-effects model of the abundance of strain Mt1B1 in feces.**

(see below)

**Table S5. p-values associated with the mixed models applied to the abundance of the 12 strains assessed from qPCR data.**

(see below)

**Table S6. Statistical analysis of the abundance of strain Mt1B1 and PFU/CFU ratios in intestinal samples.**

(see below)

**Table S7. Statistical analysis of the abundance of strain 55989 and PFU/CFU ratios in intestinal samples.**

(see below)

### Table S2. Closest phage homologs to phage P3, P10 and P17.

| Description | Max score | Total score | Query cover | E value | Identity | Accession |
|---|---|---|---|---|---|---|
| **Homologs to Phage P3 (40.1 kb)** | | | | | | |
| **Description** | **Max score** | **Total score** | **Query cover** | **E value** | **Identity** | **Accession** |
| Enterobacteria phage K1F, complete genome | 26378 | 58105 | 92% | 0.0 | 95.90% | AM084414.1 |
| Escherichia phage vB_EcoP_F, complete genome | 26092 | 59661 | 93% | 0.0 | 95.82% | KY295894.1 |
| Escherichia phage LM33_P1, complete genome | 17570 | 52746 | 87% | 0.0 | 94.66% | LT594300.1 |
| Escherichia phage PE3-1, complete genome | 17538 | 51871 | 84% | 0.0 | 94.98% | KJ748011.1 |
| Escherichia phage JSS1, complete genome | 17533 | 51844 | 84% | 0.0 | 94.98% | KX689784.2 |
| Escherichia virus Vec13, complete genome | 17457 | 52867 | 87% | 0.0 | 94.16% | MH400309.1 |
| **Homologs to Phage P10 (45.1 kb)** | | | | | | |
| **Description** | **Max score** | **Total score** | **Query cover** | **E value** | **Ident** | **Accession** |
| Escherichia virus AAPEc6, complete genome | 31124 | 60643 | 89% | 0.0 | 92.77% | KX279892.2 |
| Escherichia virus VEc3, complete genome | 29724 | 53841 | 83% | 0.0 | 92.75% | MG251390.1 |
| Escherichia phage vB_EcoP_C, complete genome | 27353 | 62103 | 90% | 0.0 | 94.08% | KY295892.1 |
| Escherichia virus mutPK1A2, complete genome | 25453 | 59969 | 90% | 0.0 | 93.81% | MG004687.1 |
| Escherichia virus K1E, complete genome | 25340 | 62944 | 89% | 0.0 | 94.36% | KY435490.1 |
| Escherichia phage vB_EcoP_KAW1A4500, complete genome | 23865 | 57685 | 88% | 0.0 | 92.75% | MK373773.1 |
| **Homologs to Phage P17 (150.9 kb)** | | | | | | |
| Description | Max score | Total score | Query cover | E value | Ident | Accession |
| Escherichia phage ESCO13, complete genome | 63247 | 2.407e+05 | 93% | 0.0 | 98.10% | KX552041.2 |
| Escherichia phage vB_EcoM-Ro121lw, complete genome | 63032 | 2.440e+05 | 94% | 0.0 | 98.28% | MH160766.1 |
| Escherichia phage vB_EcoM_Schickermooser, complete genome | 62626 | 2.396e+05 | 93% | 0.0 | 98.17% | MK373788.1 |
| Escherichia phage phAPEC8, complete genome | 45930 | 2.345e+05 | 91% | 0.0 | 97.74% | JX561091.1 |
| Escherichia phage vB_EcoM-Ro121c4YLVW, complete genome | 43886 | 2.459e+05 | 94% | 0.0 | 98.33% | MH051333.1 |
| Escherichia phage ESCO5, complete genome | 43853 | 2.332e+05 | 91% | 0.0 | 98.29% | KX664695.2 |

Magablast tool from NCBI (https://blast.ncbi.nlm.nih.gov/Blast.cgi) was used to search for the closest homologs ranked in decreasing values of query cover (Last update august 2019). Only the top 6 homologs are shown.

**Table S4. Statistical analysis of the colonization level of strain Mt1B1 in fecal samples**

ANOVA

day: testing difference between the different days

phage: effect of the phage exposure

| Response: | scale(log10(CFU.g)) | | |
|---|---|---|---|
| | Chisq | Df | Pr(>Chisq) |
| day | 15,4517 | 3 | 0,0015 |
| phage | 7,0961 | 1 | 0,0077 |
| day:phage | 12,8145 | 3 | 0,0051 |

Post-hoc tests Comparison between “phage” and “no phage” at the different time points

| | contrast | estimate | SE | df | t.ratio | p.value |
|---|---|---|---|---|---|---|
| Day=14 | no phage vs. phage | -0,20947 | 0,3581942 | 72,33 | -0,585 | 0,5605 |
| Day=15 | no phage vs. phage | 1,217633 | 0,3581942 | 72,33 | 3,399 | 0,0011 |
| Day=16 | no phage vs. phage | 0,946077 | 0,3581942 | 72,33 | 2,641 | 0,0101 |
| Day=17 | no phage vs. phage | 0,678505 | 0,3581942 | 72,33 | 1,894 | 0,0622 |

Bacterial abundance (CFU) as a function of time (day) and exposure to phages. Random effects include individual IDs as well as the cage in which they were reared. Overall analysis of variance (ANOVA) reveals significant effects of *day* (p=0.001469), *phage* (p=0.007725) and their *interaction* (p=0.005056). The *post-hoc* Tukey comparisons displayed below were performed between mice exposed to phage and not exposed within each day.

**Table S5. p-values associated with the mixed models applied to the abundance of the 12 strains assessed from qPCR data**

day 14: variations within samples day 14
day 17: variations within samples day 17
experiment: variations taking in account the 3 days (0, 14 and 17)
phage: variations caused by phage exposition

| ref# | I48 | YL44 | YL27 | YL32 |
|---|---|---|---|---|
| Strain name | *Bacteroides caecimuris* | *Akkermansia muciniphila* | *Muribaculum intestinale* | *Clostridium clostridioforme* |
| day 14 | 0,238 | 0,354 | 0,403 | 0,182 |
| day 17 | 0,45 | 0,01 | 0,838 | 0,01 |
| experiment | 0,078 | 0,862 | 0,442 | 0,179 |
| phage | 0,28 | 0,071 | 0,867 | 0,275 |

| ref# | I46 | I49 | YL58 | YL45 |
|---|---|---|---|---|
| Strain name | *Clostridium innocuum* | *Lactobacillus reuteri* | *Blautia coccoides* | *Turicimonas muris* |
| day 14 | 0 | 0,279 | 0 | 0,55 |
| day 17 | 0,782 | 0,256 | 0,022 | 0,035 |
| experiment | 0,509 | 0,556 | 0,338 | 0,295 |
| phage | 0,269 | 0,269 | 0,434 | 0,434 |

| ref# | KB1 | YL31 | YL2 | KB18 |
|---|---|---|---|---|
| Strain name | *Enterococcus faecalis* | *Flavonifractor plautii* | *Bifidobacterium longum s* | *Acutalibacter muris* |
| day 14 | 0,022 | 0,078 | NA | NA |
| day 17 | 0,269 | 0,011 | NA | NA |
| experiment | 0,112 | 0,684 | NA | NA |
| phage | 0,403 | 0,579 | NA | NA |

The model tested variations within day 14, within day 17 and also variations on the samples taking into account days 0, 14 and 17.
The model also tested variations on each strain taking into account the phage exposure.

**Table S6. Statistical analysis of the abundance of strain Mt1B1 and PFU/CFU ratios in intestinal sections**

**CFU abundance in sections**

ANOVA/ANODE

| | scale(log10(CFU.g+1)) | | | |
|---|---|---|---|---|
| | Chisq | Df | Pr(>Chisq) | |
| section | 77,354 | 4 | 6,33E-16 | *** |
| phage | 25,713 | 1 | 3,96E-07 | *** |
| section:phage | 10,113 | 4 | 0,03857 | * |

Post-hoc tests

| | Comparison between “phage” and “no phage” in the different sections | | | | | |
|---|---|---|---|---|---|---|
| | contrast | estimate | SE | df | t.ratio | p.value |
| ileum_lumen | no phage vs. phage | 1,0723044 | 0,2959727 | 110,16 | 3,623 | 0,0004 |
| ileum_mucosa | no phage vs. phage | 0,9855932 | 0,2959727 | 110,16 | 3,33 | 0,0012 |
| cecum | no phage vs. phage | 0,1633725 | 0,2959727 | 110,16 | 0,552 | 0,5821 |
| colon_lumen | no phage vs. phage | 0,525542 | 0,2959727 | 110,16 | 1,776 | 0,0786 |
| colon_mucosa | no phage vs. phage | 1,2514249 | 0,2959727 | 110,16 | 4,228 | <.0001 |

Bacterial abundance (CFU) as a function of organ and exposure to phages. Random effects include individual IDs as well as the cage in which they were reared. Overall analysis of variance (ANOVA) reveals significant effects of *organ* (p=6.33E-16), *phage* (p=3.96E-07) and their *interaction* (p=0.03857). The post-hoc Tukey comparisons displayed below were performed between the mice exposed to phage and not exposed within each day.

**PFU/CFU ratios**

ANOVA – ANODE

| Response: | scale(L.ratio) | | |
|---|---|---|---|
| | Chisq | Df | Pr(>Chisq) |
| organ | 2,4925 | 2 | 0,287582 |
| group[a] | 9,7241 | 1 | 0,001819 |
| organ:group | 0,7109 | 1 | 0,399152 |

[a] group= luminal or mucosal

Post-hoc test

| organ | contrast | estimate | SE | df | t.ratio | p.value |
|---|---|---|---|---|---|---|
| colon | lumen-mucosa | 0,5535417 | 0,3440682 | 62,13 | 1,609 | 0,1127 |
| ileum | lumen-mucosa | 0,9637997 | 0,3440682 | 62,13 | 2,801 | 0,0068 |

The ratios of phages over bacteria abundance (PFU/CFU) as a function of organ and group (mucosa or lumen). Random effects include individual IDs as well as the cage in which they were reared. Overall analysis of variance (ANOVA) reveals no significant effects of *organ* (p=0.287582), but significant of *group* (p=0.001819). The post-hoc Tukey comparisons displayed below were performed between the lumen and mucosa data. Post-hoc tests revealed a significant difference between lumen and mucosa of the ileum (p=0.0068).

**Table S7. Statistical analysis of the abundance of strain 55989 and PFU/CFU ratios in intestinal sections**

**CFU abundance in sections**

ANOVA/ANODE

| | scale(log10(CFU.g+1)) | | | |
|---|---|---|---|---|
| | Chisq | Df | Pr(>Chisq) | |
| section | 164.4484 | 4 | < 2.2e-16 | *** |
| phage | 7.2457 | 1 | 0.007107 | ** |
| section:phage | 3.2958 | 4 | 0.509598 | * |

Post-hoc tests

Comparison between "phage" and "no phage" in the different sections

| | contrast | estimate | SE | df | t.ratio | p.value |
|---|---|---|---|---|---|---|
| ileum_lumen | no phage vs. phage | 1.013 | 0.383 | 25.9 | 2.648 | 0.0136 |
| ileum_mucosa | no phage vs. phage | 0.751 | 0.383 | 25.9 | 1.962 | 0.0605 |
| colon_lumen | no phage vs. phage | 0.688 | 0.383 | 25.9 | 1.798 | 0.0839 |
| colon_mucosa | no phage vs. phage | 0.204 | 0.383 | 25.9 | 0.532 | 0.5990 |

Bacterial abundance (CFU) as a function of organ and exposure to phages. Random effects include individual IDs as well as the cage in which they were reared. Overall analysis of variance (ANOVA) reveals significant effects of *organ* (p<2.2E-16), *phage* (p=0.007) but not their *interaction* (p=0.509). The post-hoc Tukey comparisons displayed below were performed between the mice exposed to phage and not exposed within each day.

**PFU/CFU ratios**

ANOVA – ANODE

| | | | | |
|---|---|---|---|---|
| Response: | scale(L.ratio) | | | |
| | Chisq | Df | Pr(>Chisq) | |
| organ | 18.5313 | 1 | 1.671e-05 | *** |
| group[a] | 1.3823 | 1 | 0.2397 | |
| organ:group | 1.2474 | 1 | 0.2641 | |

[a] group= luminal or mucosal

Post-hoc test

| organ | contrast | estimate | SE | df | t.ratio | p.value |
|---|---|---|---|---|---|---|
| colon | lumen-mucosa | 0.872 | 0.373 | 31.1 | 2.337 | 0.0261 |
| ileum | lumen-mucosa | 1.475 | 0.373 | 31.1 | 3.784 | 0.0007 |

The ratios of phages over bacteria abundance (PFU/CFU) as a function of organ and group (mucosa or lumen). Random effects include individual IDs as well as the cage in which they were reared. Overall analysis of variance (ANOVA) reveals a significant effect of *organ* (p=1,671E-5), but not significant of *group* (p=0.2397). The post-hoc Tukey comparisons displayed below were performed between the lumen and mucosa data. Post-hoc tests revealed significant differences between lumen and mucosa for both the *colon* (p=0.0261) and the *ileum* (p=0.0007).
